# Deep coalescent history of the hominin lineage

**DOI:** 10.1101/2024.10.17.618932

**Authors:** Trevor Cousins, Richard Durbin, Regev Schweiger

**Affiliations:** Department of Genetics, University of Cambridge, Cambridge, UK; Department of Human Genetics and Computational Medicine, Gray Faculty of Medical and Health Sciences, Tel Aviv University, Tel Aviv, Israel; Blavatnik School of Computer Science and AI, Tel Aviv University, Tel Aviv, Israel

## Abstract

Coalescent-based methods are widely used to infer population size histories, but existing analyses have limited resolution at deep time scales (*>*2 million years ago). Here, using new high-quality telomere-to-telomere genome assemblies, we extend the scope of such inference and reanalyse an ancient peak in human effective population size around 3–6 million years ago, showing that coalescent-based inference can reach much further into the past than previously thought. Furthermore, peaks at approximately the same time as in humans are also observed in chimpanzee and bonobo, but not in gorilla or orangutans. We show that this pattern is unlikely to be an artefact of model violations and discuss its potential implications for understanding hominin evolutionary history. It is known that such peaks can arise from ancestral genetic structure, and consequently we suggest that a parsimonious explanation for the contemporaneous peaks in humans, chimpanzees and bonobos would be a period of complex speciation before final separation of *Homo* from *Pan*, in contrast to a scenario where these lineages diverge cleanly from a panmictic common ancestor.

## 1 Introduction

The study of human population size over time provides a valuable lens through which to explore key questions about our evolutionary past [1]. Effective population size history, *N* (*t*), taken here as half of the inverse of the coalescence rate at time *t*, can be inferred from present-day genome sequences by examining patterns of genetic variation within and between populations. Various methods have been developed for such inference, using information such as the site frequency spectrum [2, 3], linkage disequilibrium [4, 5], or the genomic distribution of heterozygosity [6, 7, 8]. These approaches have provided important insights into the evolutionary history of humans, including the magnitude of exponential growth over the last 20 thousand years (kya), the timing and severity of the out-of-Africa bottleneck around 50-60 kya, and divergence times among human subpopulations as well as between humans, Neanderthals and Denisovans [9, 10]. However, methods to infer human *N* (*t*) have typically focused on evolution within the past 1–2 million years (Mya), even though the mean coalescence time in humans is ∼1 Mya and present-day genomes should therefore contain information about substantially older events. Extending inference to these periods could help illuminate the emergence of the genera *Homo* (∼3 Mya [11, 12]) and *Australopithecus* (∼4.5 Mya [13]), as well as the ancestral population of humans, chimpanzees, and bonobos (∼6 Mya [14]), providing insight into deep hominin evolutionary history.

One class of approaches for studying these timescales analyses divergence among species by comparing genome sequences from humans, chimpanzees, and gorillas. Methods such as CoalHMM [15] and its successors [14, 16] employ coalescent hidden Markov models (HMM) in which the hidden states represent the ordering of coalescent events, and in some formulations also ancestral coalescence times. While these methods typically estimate divergence times and ancestral population sizes, they focus on a limited set of parameters, rather than reconstructing a continuous population size history through time. A separate class of methods infers population size histories within species, most notably the pairwise sequentially Markovian coalescent (PSMC) method [6] and its successors [7, 17, 18, 19, 20]. These methods use a hidden Markov model in which hidden states represent discretised local coalescence-time intervals whose transition probabilities are derived from the sequentially Markovian coalescent (SMC) model [21] or its variant SMC’ [22], which includes an additional class of coalescent events missed in the original SMC. In principle, estimates of *N* (*t*) from these methods may extend beyond 1-2 Mya, but inference at such depths has typically not been discussed, partly because the robustness of the SMC approximation under model violations at these timescales remains unclear. As a result, population dynamics during deep hominin evolutionary history remain poorly understood.

Another key limitation in inferring deep hominin evolutionary history has been the quality of available genome assemblies. Most human demographic analyses have relied on short-read data aligned to reference genomes derived from incomplete assemblies that collapse or omit difficult genomic regions [15, 23, 24, 14, 16]. In addition, assemblies for non-human apes are typically aligned to the human reference genome, or incorporated into multiple genome alignments, despite millions of years of divergence between these species. This evolutionary distance can introduce reference bias in variant calling [25], which may in turn affect down-stream demographic inference. Recently, the advent of accurate long-read sequencing [26, 27] has enabled telomere-to-telomere (T2T) assemblies that provide highly accurate, haplotype-resolved reference genomes for humans [27, 28, 29] as well as for a chimpanzee, bonobo, gorilla, Bornean orangutan, and Sumatran orangutan [30]. These assemblies expand the fraction of the genome that can be analysed while also substantially reducing sequencing and assembly errors. Because demographic inference is highly sensitive to false mutation calls and missing sequence, the increased completeness and accuracy of T2T genomes should enable more reliable inference at deeper evolutionary timescales.

Here we extend coalescent-based population size inference further into deep timescales. We examine an ancient peak in inferred effective population size around 3–6 Mya observed in PSMC-type analyses of human, chimpanzee, and bonobo genomes, but not in gorilla or orangutan genomes, a signal whose traces have appeared in several earlier studies but has received little attention [6, 9, 10, 24, 31, 32]. Notably, this interval overlaps with the estimated divergence of the *Homo* and *Pan* lineages (henceforth the *Homo–Pan* (HP) divergence, ∼6 Mya [14, 16, 30]), a transition debated as either being a clean split or having a period of post-divergence gene flow [33, 34, 14, 35, 36, 37]. Using T2T genome assemblies, we reanalyse this signal and assess potential model violations affecting these inferences. Our results are consistent with gene exchange between the ancestral human and chimpanzee–bonobo lineages prior to their final separation, highlighting the potential of coalescent-based inference to provide new insight into deep hominin evolutionary history.

## 2 Results

### 2.1 An ancient peak in human, chimpanzee, and bonobo population size history

We used the T2T haplotype-resolved HG002 human genome reference [27, 28, 29] to align the alternate assembly to the primary assembly and call variants (see Methods). We additionally aligned 26 high-coverage genomes from various populations from the 1000 Genomes Project (1000GP) [38, 39, 32, 40] to the HG002 primary assembly and performed variant calling (see Methods). For each of these genomes, we used the pairwise SMC implementation in MSMC2 [41], which we note includes the SMC’ correction and we refer to from now on as PSMC’, to infer an effective population size (*N* (*t*)) curve over time. Relative to previous similar analyses, we extended the inference by applying more stringent variant calling and by fitting the model with more time bins and iterations, and fewer parameter constraints [42] (see Methods), allowing greater resolution at timescales beyond 2 Mya. The resulting *N* (*t*) estimates span from 10 kya to 10 Mya, with consistent bootstrap results between 20 kya and 8 Mya (Figure 1a), extending substantially deeper into the past than has been achieved in previous studies [6, 9, 10, 31]. The inference reveals two peaks, one approximately 400 kya and a second older one, starting around 3 Mya and extending further back in time, reaching a maximum around 4.5 Mya with a population size of ∼32,500, then decreasing until around 6 Mya (shaded in yellow in Figure 1). This pattern is stable across 20 bootstrap replicates and consistent across each of the 26 1000GP populations (Figure A1).

**Figure 1.**
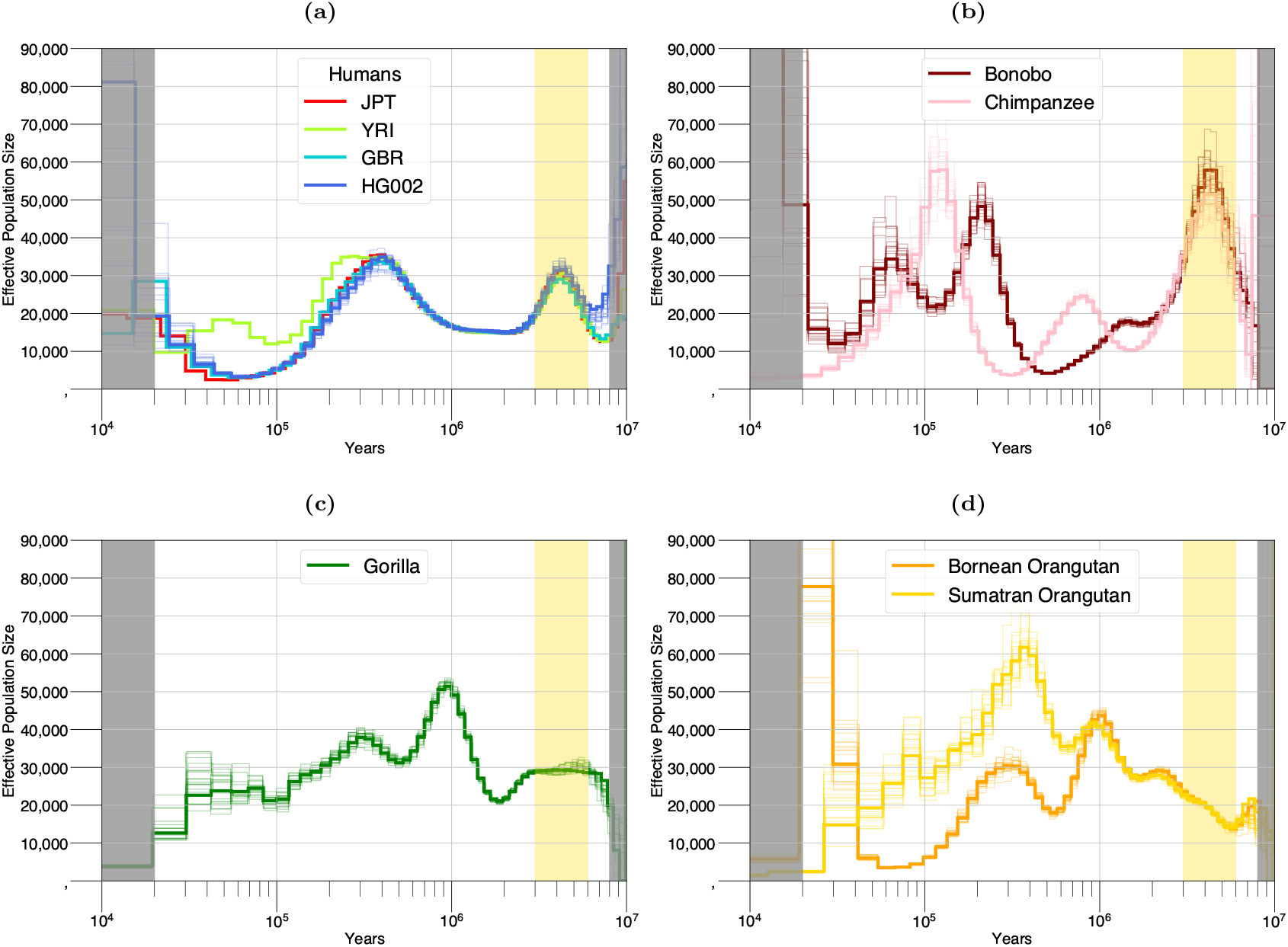
Inference from PSMC’ in humans, chimpanzees and bonobos, gorillas, and orangutans. **(a)** Estimates of population size history in humans, using HG002 and 3 individuals each from distinct populations in the 1000 Genomes project. **(b)** Estimates of population size history in chimpanzee and bonobo, using mPanTro3 and mPanPan1. **(c)** Estimates of population size history in gorilla, using mGorGor1. **(d)** Estimates of the population size history in Bornean and Sumatran orangutans, using mPonAbe1 and mPonPyg2, respectively. The faint lines in each plot show 20 iterations of non-parametric block bootstrapping.

As the ancient peak occurs around the estimated time at which humans split from the common ancestor of chimpanzees and bonobos, it is natural to ask if a similar signal exists in chimpanzee and bonobo population size history, or in other great apes in general. Using T2T assemblies from the T2T-Primates project [30], we therefore applied PSMC’ to chimpanzee (mPanTro3), bonobo (mPanPan1), gorilla (mGorGor1), Sumatran orangutan (mPonAbe1) and Bornean orangutan (mPonPyg2), to infer their population size histories. First, we recovered an ancient peak in the chimpanzee and bonobo, occurring at the same time as the peak seen in the human analysis (Figure 1b). As with humans, this extends further back in time a signal present in previous analyses but not discussed [24]. We see the ancient peak more clearly than previously because we avoid grouping old time intervals and due to the lower error rates of the T2T assemblies, as for human. The fact that there are peaks in all three of humans, chimpanzee and bonobo that occur within the time frame of the estimated HP divergence prompts the question of whether these peaks signify shared demographic events before the lineages fully separated into distinct species. We note that the effective population size of the chimpanzee and bonobo ancient peak is significantly higher than that of humans; we show in Figure A2 that such height differences can occur from shared ancestral structure where gene flow rates are asymmetric, while acknowledging that there are many other possible explanations for such shared patterns - we return to this in the Discussion.

For the gorilla (Figure 1c) and the orangutans (Figure 1d), we do not detect a discernible peak in the region between 3-6 Mya. This is consistent with the fact that gorillas and orangutans genetically separated from humans, chimpanzees and bonobos prior to this period, and so would not be expected to share their history at that time (gorillas are estimated to have separated from the ancestral human, chimpanzee and bonobo population around 10 Mya, and their joint ancestral population is estimated to have separated from orangutans around 19 Mya [30]). Moreover, it supports the claim that the ancient peak observed in humans, chimpanzees and bonobos is not an artefact of the method or of the data processing, as that would result in a peak with gorillas and orangutans as well.

### 2.2 The fraction of uncoalesced genome as a function of time

The power of SMC based methods to infer *N* (*t*) at ∼3-6 Mya relies on there being sufficiently many places in the genome where the two input lineages coalesce more anciently than this time. To assess whether this is reasonable to expect, we note that under a simple panmictic model the observed range of heterozygosity in humans (e.g., ∼0.00069 for PEL to ∼0.001 for MSL in 1000GP) would correspond to a constant effective diploid population size of 13,800-20,000, using a mutation rate of 1.25e-8 per basepair per generation. Given an exponential distribution of coalescence times, and a generation time of 29 years, we expect a mean coalescence time of 800,400-1,160,000 years, and that the coalescence time will be older than 5 Mya in 0.02%-1.3% of the genome. Similarly, given the observed heterozygosity in the chimpanzee sample (∼0.0008 for mPanTro3) and a mutation rate of 1.78e-8 per base pair per generation and generation time of 24, we expect a mean coalescence time of 540,000 years and that the coalescence time will be older than 5 Mya in 0.01% of the genome.

As more direct confirmation from the data, we inferred the fraction of the genome that has not yet coalesced as a function of time using PSMC’ decoding (see Methods). At least 1% of the human, gorilla, and orangutan genome is estimated not to have coalesced by 5 Mya, with approximately 0.5% remaining in the chimpanzee and bonobo genome at this time (Figure 2a). All of the non-human primates are estimated to have fully coalesced between 7.4 Mya and 11 Mya. Notably, however, even at 10 Mya an average of 0.06% of the human genome remains uncoalesced, with full coalescence not being reached until beyond 12.5 Mya in the YRI and HG002 samples.

**Figure 2.**
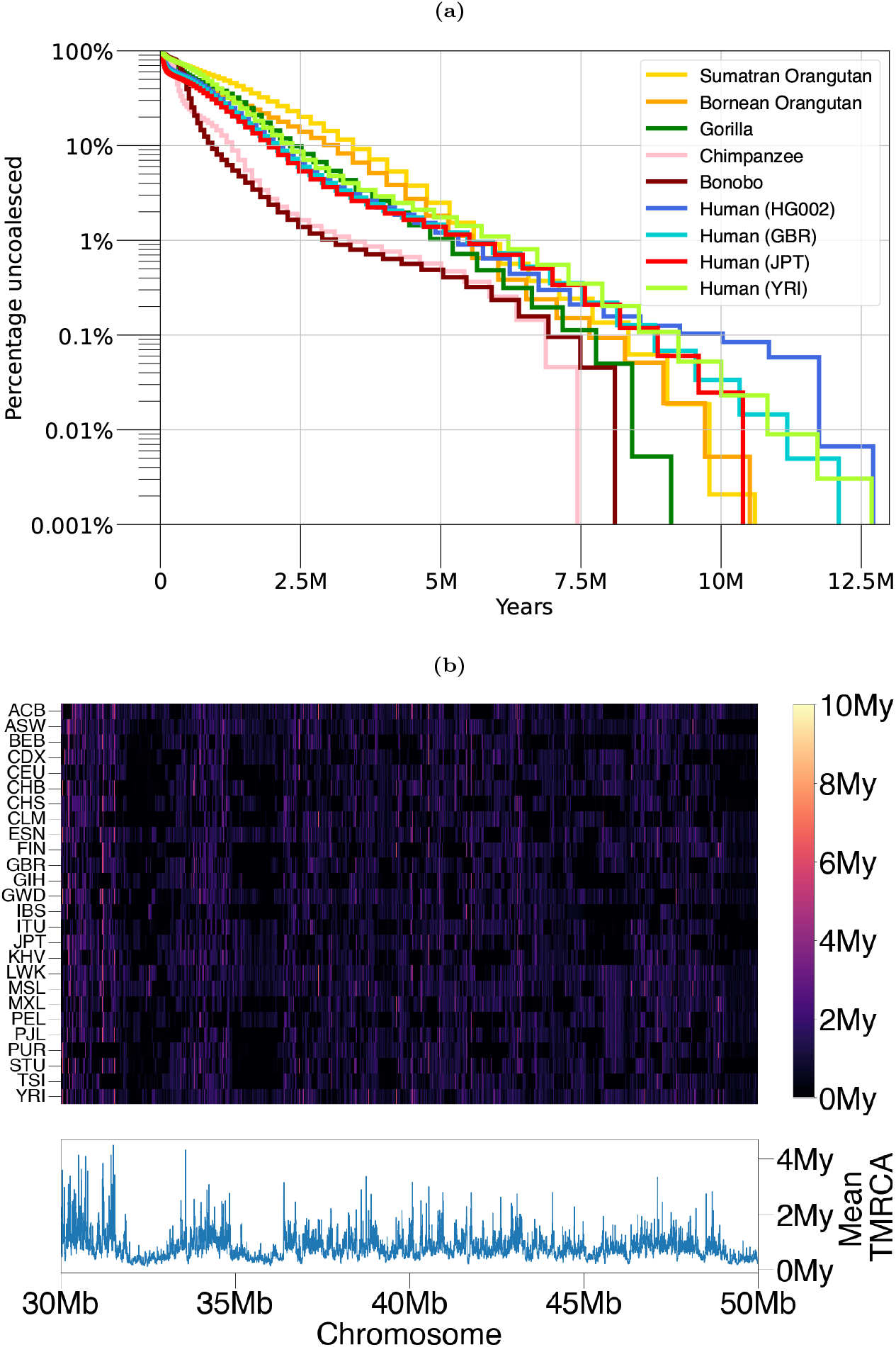
**(a)** Inferred fraction of uncoalesced genome as a function of time, for each sample from Figure 1. **(b)** (Top) Posterior mean TMRCAs from 26 individuals, each from a distinct population in the 1000 Genomes project, for 20Mb on chromosome 20. The bottom panel shows the mean of the top panel.

In humans from the 1000GP, ancient regions tend to be shared among individuals (Figure 2b); e.g. positions with an inferred posterior mean TMRCA of over 3 Mya show an average correlation of *r*^2^=0.21 across individuals (Figure A3). We note that further back in time, segments of shared ancestry become increasingly shorter because of longer exposure to recombination, and that these are scattered across the genome. For example we estimate that African individuals harbor roughly 4,000 non-contiguous segments whose coalescence time is older than 4 Mya, and roughly 400 segments whose coalescence time is older than 6 Mya (Table 1; see Methods). This number of distinct segments is sufficient to give robust coalescence estimates, consistent with the bootstrap results. We note that previous simulation studies [43, 44] and theoretical analysis [45] have shown that posterior mean estimates of true ancient coalescent times substantially older than the mean tend to be significantly downward biased, which may indicate an even older age for detected ancient regions. We repeated the analysis using an independent method, Relate [46], which does not rely on the assumptions of the PSMC’ model and uses multiple sequences jointly; this also identifies sufficient coalescence material exists in this time period (see Methods and Figure A4).

**Table 1.** The number of ancient non-contiguous segments inferred in different African populations from the 1000GP project. We count segments by looking for regions where the posterior probability of coalescence as old or older than the time given in each column is greater than 0.9.

| Population | Sample | 1 Mya | 2 Mya | 3 Mya | 4 Mya | 5 Mya | 6 Mya |
| --- | --- | --- | --- | --- | --- | --- | --- |
| ESN | HG03515 | 54,169 | 17,213 | 7,811 | 3,952 | 1,841 | 472 |
| YRI | NA18488 | 54,486 | 17,229 | 7,668 | 3,841 | 1,752 | 426 |
| MSL | HG03212 | 53,817 | 17,243 | 7,661 | 3,910 | 1,947 | 494 |
| GWD | HG02568 | 54,493 | 16,893 | 7,358 | 3,667 | 1,721 | 397 |
| ACB | HG01882 | 55,368 | 17,590 | 7,883 | 4,129 | 1,909 | 457 |
| ASW | NA19625 | 54,974 | 17,395 | 7,798 | 3,984 | 1,852 | 439 |
| LWK | NA19017 | 54,644 | 17,709 | 8,105 | 4,050 | 1,874 | 496 |

### 2.3 Model violations and their effects

As with all statistical frameworks, the PSMC’ model that we use relies on specific assumptions that may not hold true in reality. Deviations from these assumptions could have varying impacts on inference across different temporal scales, and could potentially account for some or all of the observed ancient peak. We therefore considered how departures from the modelling assumptions could plausibly affect inference of population size history in the deep past.

To this end, we considered the following possible issues. First, repeats may manifest as an artifactual heterozygous signal, leading to long segments with high heterozygosity, which then leads to an apparent excess of ancient coalescence events. Second, variation in mutation rates across different genomic regions could result in some regions having a higher mutation density, which might be interpreted as an older TMRCA, while others with lower mutation density might be interpreted as having a younger TMRCA. This can lead to systematic biases in the inference of accurate evolutionary histories. Third, the human mutation rate is estimated to have slowed down over time [47, 48, 49, 50], while in the PSMC’ model that we use it is assumed constant. Fourth, variability in recombination rates across the genome (in particular at recombination hotspots) is also a departure from model assumptions [51, 52, 53, 54, 55]. Fifth, at longer timescales there may become a significant probability of recurrent mutations at the same site, which are not modelled in PSMC’. Sixth, background selection, which has been estimated to be pervasive in humans [56, 57], has been shown to distort the inference of the coalescence rate [58, 59, 60]. Finally, balancing selection can also manifest as an excess in ancient coalescence events [61].

We conducted an extensive set of simulation studies to analyse how these violations may affect inference, which we describe fully in the Appendix. We concluded that it is unlikely any of these issues or violations could cause a false ancient peak of the type we see in real data. Notably, we observed that if there are large variations in the mutation rate across the genome, then false inflations of inferred *N* (*t*) are observed. However, the degree of rate variation required to generate this is larger than the degree of this inferred in humans, and moreover their ancient inferred *N* (*t*) has high variance, contrasting the low variance we have shown in humans in the ∼3-6 Mya peak.

## 3 Discussion

We have demonstrated the applicability of coalescent-based inference to examine deep evolutionary history in the hominin lineage. Specifically, we highlighted and extended the analysis of a peak in ancient population size history inferred by the PSMC’ model, which is reliably estimated from human, chimpanzee and bonobo data, but not in gorilla or orangutan data. The human, chimpanzee and bonobo peaks look qualitatively similar and span approximately the same time period, suggesting that they may reflect the same event prior to, or during, *Homo-Pan* (HP) speciation.

We discussed several ways in which the data may violate the model underlying our analysis, and concluded that none are likely to generate a false ancient peak of the type seen. Nevertheless, it may be useful to incorporate some of these as extensions to the model, modifying the transition and emission probabilities within the underlying HMM accordingly. Indeed, some work has already been done towards this goal [62, 63], and future modification could further address slowdown or genomic variability in mutation rate, recombination hotspots, and/or selection, so as to better understand demographic change and speciation across a variety of species.

A straightforward interpretation of the shared ancient peak in humans, chimpanzees and bonobos is that it reflects true changes in population size. Indeed, for humans, changes in inferred *N* (*t*) since ∼80 kya have been associated with the out-of-Africa event followed by more recent recent population size increases [7, 19]. The ancient peak may similarly reflect ancient hominin population expansion and decline in the late Miocene and the Pliocene. To the extent that the same ancient peak exists in chimpanzees and bonobos, it may reflect common environmental conditions affecting similar evolutionary processes in the different species. Even if the ancestral population were structured, shared environmental changes may have contributed to changes in inferred *N* (*t*).

However, there are other possible explanations. The separation and reconnection of local populations alters the amount of possible gene flow between them, which affects the inferred coalescence rate [64, 65, 66, 67, 68]. In a structured population (defined as a population whose evolutionary history was not continuously panmictic) the coalescence rate decreases relative to a panmictic population of the same size, thus the effective population size (taken throughout this paper as half of the inverse of the coalescence rate) increases. Indeed, in the original PSMC publication Li and Durbin demonstrate that a pulse admixture event can generate a peak in effective population size inference [6], and in a more thorough analysis Mazet et al. show that the inferred human effective population size as inferred by PSMC and related methods - including the ancient peak - can be equally well obtained by a set of populations with varying migration rates between them without any changes in size [67]. Therefore, it is possible that the ancient peak is a signature of ancient population structure in some form.

One such possible structured event may be complex speciation between the HP lineages, in which there was some form of gene flow between the *Homo* and *Pan* lineages after their initial separation and prior to their final loss of ability to exchange genetic material. Previous publications have put forward evidence for this type of model [33, 34, 14, 35], although other analysis has favoured a model without any gene flow [36, 37]. Broadly, these studies employ two types of models when testing for historical structure. One class of model has a panmictic ancestral population cleanly split into two distinct populations that later exchange genes in a pulse gene flow event. The other assumes that after initial divergence there is some level of continuous gene flow before full separation. Innan and Watanabe proposed there was no strong support for continuous gene flow after initial divergence, and that a clean split best fit the data [36]. This result was reinforced by Yamamichi et al., whose maximum likelihood pulse model also favoured a clean split [37] (we note that their pulse model was slightly different in that after a period of isolation the two populations merged again into a panmictic population before later cleanly separating). Patterson et al. proposed that a pulse gene flow event did occur, because of differences in the estimated divergence time between the autosomes and X chromosome [33], although this approach has been criticised [69, 70, 37]; specifically it was argued that expected coalescent variance and lineage-specific differences in male mutational bias driven mutation could provide an alternative explanation for the observed signal [70]. Using a likelihood ratio test considering divergence times across the genome, Yang rejected the null hypothesis of an absence of gene flow (noting that their model is compatible with either a pulse or continuous gene flow scenario) [34]. With an HMM that explicitly tests for immediate or prolonged separation with a period of continuous gene flow, Mailund et al. proposed that a model with an extended period of gene flow better fits the data [14]. Galtier introduced Aphid, which distinguishes between discordant coalescent trees (ones that do not match the species trees) generated by ILS or gene flow [35]. Aphid leverages the fact that multispecies coalescent trees affected by gene flow tend to have shorter branches, and coalescent trees affected by incomplete lineage sorting longer branches, than the average coalescent tree. This information provided strong support for a model in which there was ancient gene flow between human, chimpanzee, and even gorilla, after the initial human-chimpanzee split (in this model the flow was via a pulse migration event).

We propose that a parsimonious explanation for the shared ancient peak in humans and chimpanzee-bonobo is that there was substantial shared genetic population structure in their ancestry around the time of HP separation, i.e. that this reflects some form of complex speciation. We do not provide an explicit model for this structure, e.g. pulse versus continuous, or a specific number of demes with specific gene flows, because the experience of the field is that such models are underconstrained. However we contend that there is qualitative evidence to support this proposal, independent of the simple suspicious coincidence of timing. Notably, CoalHMM [71] and TRAILS [16], which assume a panmictic ancestral human-chimpanzee-bonobo population and a clean split, estimate the ancestral effective population size as 177,368 and 167,400, respectively. Both of these estimates are not well aligned with the PSMC’ inference in this period (Figure 1), and are more than 10 times the long-term effective population size of humans and chimpanzees (defining “long-term effective population size” as the value inferred from the observed heterozygosity using the equation 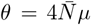. In humans 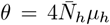 is roughly 0.0008, which assuming *µ*_*h*_=1.25e-8 gives human long-term effective size 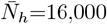 in chimpanzees, *θ* is roughly 0.001, which assuming *µ*_*c*_=1.78e-8 gives chimpanzee long-term effective size 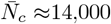). A structured ancestral population could explain the large CoalHMM and TRAILS estimates, as falsely assuming panmixia can inflate estimates of population size [67]. Moreover, Aphid [35], which provides strong support for human-chimpanzee-gorilla gene flow, estimates the size of the ancestral human-chimpanzee population to be of similar size to present-day human, chimpanzee and bonobo populations.

We note that there are believed to be other speciation/separation events in the human lineage for which we see no peak in analyses based on the PSMC model, for example divergences among early Homo lineages at 2-3 Mya, including those associated with *Homo habilis*. However, even if there was gene flow associated with these separations, a divergence event with gene flow need not always generate a peak in such analyses. In a recent study, Cousins et al. simulated various structured models in which an ancestral population diverges into two distinct populations that later exchange genes via a pulse migration event [68]. Analyses of these simulations, which were based on the same PSMC’ implementation within MSMC2 that we use here, found that the magnitude of the inferred peak is a function of the amount of admixture, the length of time between separation and reconnection, as well as the density of coalescence events during the structured period (Supplementary Figure 19 in [68]). In particular, with a human-inspired effective population size and mutation rate, if the length of time between the two admixing populations is less than 1 million years and the amount of admixture is less than 5% then there is no noticeable signal in the analysis. As an illustrative example of this effect in real data, consider the Neanderthal to non-African modern humans introgression: the amount of introgression from Neanderthals into non-Africans is estimated to have been on the order of 2% and to have occurred roughly ∼50 kya [10, 31], with the two populations having diverged ∼550kya. There is not a discernible population size peak associated with this gene flow event; rather, in humans the non-African peak around 300-400 kya in Figure 1 is shared with Africans who are not expected to have had any Neanderthal admixture - that peak is better explained by deep structure in the hominin lineage [72, 68]. Conversely, however, Schur et al. propose that the Neanderthal introgression does subtly perturb the magnitude of the peak at ∼300 kya, though they do not propose the peak is a result of this event [73].

Divergence with gene flow events not necessarily generating inferred population size peaks also plausibly explains why other reported admixtures do not have peaks associated with their reported timing. For example, de Manuel et al. estimate that 1% of the central chimpanzee genome descends from an introgression event less than 500 kya from a bonobo-related lineage [74]. Kuhlwilm et al. estimated that 0.9-4.8% of the bonobo’s genome descends from an unsampled lineage that separated ∼3 Mya and introgressed ∼600 kya [75]. Nater et al. estimated that the Sumatran orangutan continously exchanged genes with the Tapanuli orangutan around 100 kya after their separation ∼3 Mya [76]. Pawar et al. estimated that the eastern gorillas derive ∼3% of their genome from an unsampled lineage that split from a common ancestor more than 3 Mya [77]. These estimated admixture events are of relatively small magnitude and generally do not produce discernible inferred peaks at the time of introgression, reinforcing that only sufficiently strong or prolonged gene flow is expected to generate detectable features in population size trajectories.

Mazet et al. [67], as well as Tournebize and Chikhi [78], have demonstrated that PSMC inference on an individual descended from a structured stem population (a set of sub-populations that are exchanging genes at some rate) can show peaks around the periods associated with gene flow [67, 78]. In light of this framework, the peaks observed in human, chimpanzee, and bonobo population size trajectories between roughly 3–6 Mya are consistent with descent from a structured ancestral population around that time, which is around the time of human-chimpanzee lineage separation. Assuming the eventual *Homo* and *Pan* lineages descended from different subpopulations of that structured ancestry, this scenario corresponds to our suggestion of complex speciation, in which the two lineages did not split instantaneously from a single panmictic population but instead emerged from different components of a structured ancestral population connected by migration, before complete reproductive isolation was established. In the long term, we can look forward to a better resolution of what happened in this important time for human evolution based on a combination of improved genetic and fossil evidence.

## Acknowledgements

We thank Aylwyn Scally for constructive comments. A preprint version of this article has been peer-reviewed and recommended by PCI Math Comp Biol. We thank Alan Rogers, Olivier Mazet, and an anonymous reviewer for their reviews. T.C. was funded by a Wellcome Postgraduate Studentship 108864/B/15/Z, and R.D. and R.S. by Wellcome Investigator Award 207492/Z/17/Z. For the purpose of open access, the author has applied a CC BY public copyright license to any Author Accepted Manuscript version arising from this submission.

## 4 Methods

### 4.1 Processing data

We downloaded the HG002 fasta assemblies from Hansen et al. [79]. We downloaded the chimpanzee (mPanTro3), bonobo (mPanPan1), gorilla (mGorGor1), Bornean orangutan (mPonPyg2) and Sumatran orangutan (mPonAbe1) diploid fasta assemblies from Yoo et al. [30]. For each species, we processed the data through the following pipeline:

1. We used dipcall [80] with default parameters to align the alternate assembly to the primary, call variants, and designate regions of poor quality which were subsequently masked.
2. We masked all indels as well as 5 base pairs upstream and downstream of each.
3. We detected short tandem repeats using Tandem Repeat Finder with default parameters [81] and masked these.
4. We detected regions of low complexity using sdust [82] with *w* = 64 and *t* = 20 and masked these.
5. To account for multi-nucleotide mutation events [83, 84] we masked SNPs if they were within 20 base pairs of another SNP.

We took high-coverage whole-genome-sequence CRAM files for one individual in each of the 26 populations from the 1000 Genomes project [40]. We randomly chose the following individuals from each population: ACB-HG01882, ASW-NA19625, BEB-HG03006, CDX-HG02373, CEU-NA12718, CHB-NA18530, CHS-HG00443, CLM-HG01250, ESN-HG03515, FIN-HG00266, GBR-HG00118, GIH-NA20845, GWD-HG02568, IBS-HG01783, ITU-HG03977, JPT-NA18939, KHV-HG02113, LWK-NA19017, MSL-HG03212, MXL-NA19648, PEL-HG02285, PJL-HG03234, PUR-HG01171, STU-HG03753, TSI-NA20752, YRI-NA18488. We realigned these to the primary HG002 assembly using minimap2 [85]. The CRAM files were converted to BAM and indexed with samtools [86, 87]. The genotype likelihoods were calculated with bcftools mpileup [88] by skipping alignments with mapping quality less than 20 and setting the coefficient for downgrading mapping quality to 50. All indels were excluded. Variants were then designated as uncallable if the minimum mapping quality was less than 20, the minimum consensus quality was less than 20, or the coverage was less than half or more than double the mean coverage. Finally, we masked low complexity regions as detected from sdust as well as short tandem repeats as detected with Tandem Repeat Finder, as above for the genome assemblies. Uncallable positions were labelled as missing data in the input to PSMC’.

### 4.2 Mutation rates and generation times for humans and primates

In humans we set the generation time as 29 years [89] and the mutation rate per generation per base pair as *µ*=1.25e-8 [90, 91, 92, 93, 94, 95, 96]. The rates we set for the other great apes are taken from [47, 50]. In chimpanzees and bonobos we set *µ*=1.78e-8 and generation time equal to 24 years. For Gorillas we set *µ*=1.42e-8 and generation time equal to 19 years. For Orangutans we set *µ*=2.03e-8 and generation time equal to 27 years.

### 4.3 PSMC’ analysis

In all of our PSMC-based analyses, we used the implementation of the pairwise SMC’ [22] model in MSMC2 [41], which we term PSMC’. We used MSMC2 rather than the original PSMC software [6] because the latter uses a less accurate approximation to the coalescent with recombination [21], as well as binning the sequence into chunks of size 100 base pairs which reduces resolution in ancient time. We used 64 time intervals and enabled the parameters for the effective population size in each to be inferred freely. This contrasts with the default settings of previous implementations of the algorithm that force adjacent intervals to be the same, which has been shown to lead to fitting problems [42]. For the initial value for the ratio of mutation rate to recombination rate we used the default value of 4, and the recombination rate was set to be inferred as part of the EM algorithm, which was iterated for 40 iterations. For all analyses we discretised time into *D*=64 non-overlapping intervals, where the *i*-th boundary is defined by

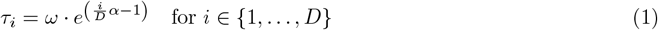

where 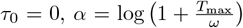, and *ω* and *T*_max_ are parameters that control the spread of time interval boundaries that are determined by the PSMC’ model.

### 4.4 Decoding of coalescence times

Using the inferred effective population size parameters, scaled recombination rate, and scaled mutation rate (*θ*), we used the PSMC’ decoding to get a posterior probability of coalescence at every position. To calculate the amount of uncoalesced genome over time, we averaged the posterior probability distributions across all positions and calculated the empirical survival function (Figure 2). The model estimates time in coalescent units (Equation (1)), where one unit corresponds to 2*N* generations for a diploid population. To convert these times to years, we first estimate long-term effective population size 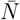 from the estimated scaled mutation rate using 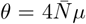, giving 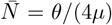, which uses the unscaled mutation rate *µ*. We then convert each inferred time point *t* by multiplying by 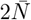 to obtain time in generations, and finally multiply by the generation time *g* (in years) to obtain time in years. Equivalently, substituting for 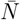, this yields 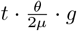. To get a point estimate from PSMC’ (Figure 2b), we take the posterior mean at each position using the midpoint of the time interval boundaries.

To calculate the number of ancient non-contiguous segments inferred in different African populations in the 1000GP (Table 1), we looked for regions where the posterior probability of coalescence older than 2 Mya, 3 Mya or 4 Mya, respectively, is greater than 0.9. We then only counted non-contiguous segments, which are all segments that are not adjacent.

### 4.5 Non-parametric bootstrapping

We can resample the data with replacement to get an estimate of the uncertainty of the parameters inferred by PSMC’. For a given individual, for each chromosome, we break the sequence up into segments of 5Mb, then reconstruct a new sequence by uniformly sampling with replacement from all 5Mb segments. The sampled segments are then concatenated together to form a contiguous sequence, on which PSMC’ inference is performed using 40 iterations. This procedure is repeated 20 times for each T2T sample in Figure 1.

### 4.6 Relate

Relate [46] is a method that estimates a genome wide ancestral recombination graph for numerous sequences. Rather than using the SMC to probabilistically infer genealogies, for each position it builds a distance matrix to describe the level of similarity between sequences, from which a rooted binary tree is constructed. The branch lengths of these trees are then estimated using an MCMC algorithm. We ran Relate on all samples in the 1000GP for GBR, CHB, and YRI. We randomly selected 10 diploid sequences from each population. To calculate the fraction of uncoalesced genome we discretised time into intervals of 500 generations and computed a histogram based on Relate’s ARG. The inferred fraction of uncoalesced genome at 5 Mya was roughly 0.3%, for each sample in each population.

### 4.7 Simulation of ancient asymmetric gene flow

Using msprime [97], we simulated 2Gb of sequence for a human and chimpanzee according to the following demographic model. A panmictic ancestral population of size 20,000 splits into proto1 and proto2 at time 6 Mya, each of size 20,000. At 3 Mya, human forms as a 80:20% mix of proto1 and proto2, respectively, and chimpanzee forms as a 50:50% mix of them. Both human and chimpanzee are of constant size 20,000 from 3 Mya to present. We used a generation time of 30 years and mutation rate of 1.25e-08. We ran PSMC’ on these simulated sequences as described in Section 4.3.

### 4.8 Reproducibility

Key scripts that were used for analysis have been uploaded to https://github.com/trevorcousins/twin_peaks.

## 5 Appendix

### 5.1 Figures

**Figure A1.**
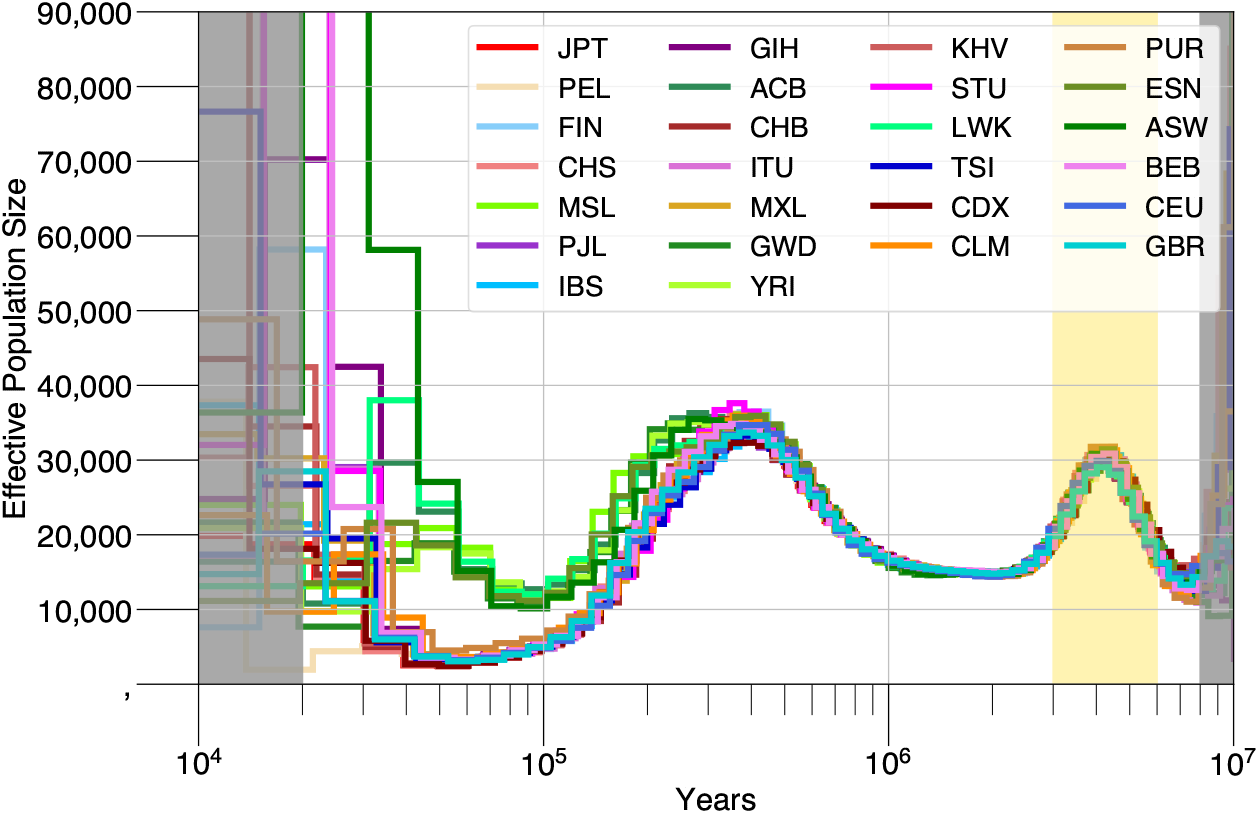
Inference from PSMC’ on 26 individuals from the 1000 Genomes project, each from a distinct population.

**Figure A2.**
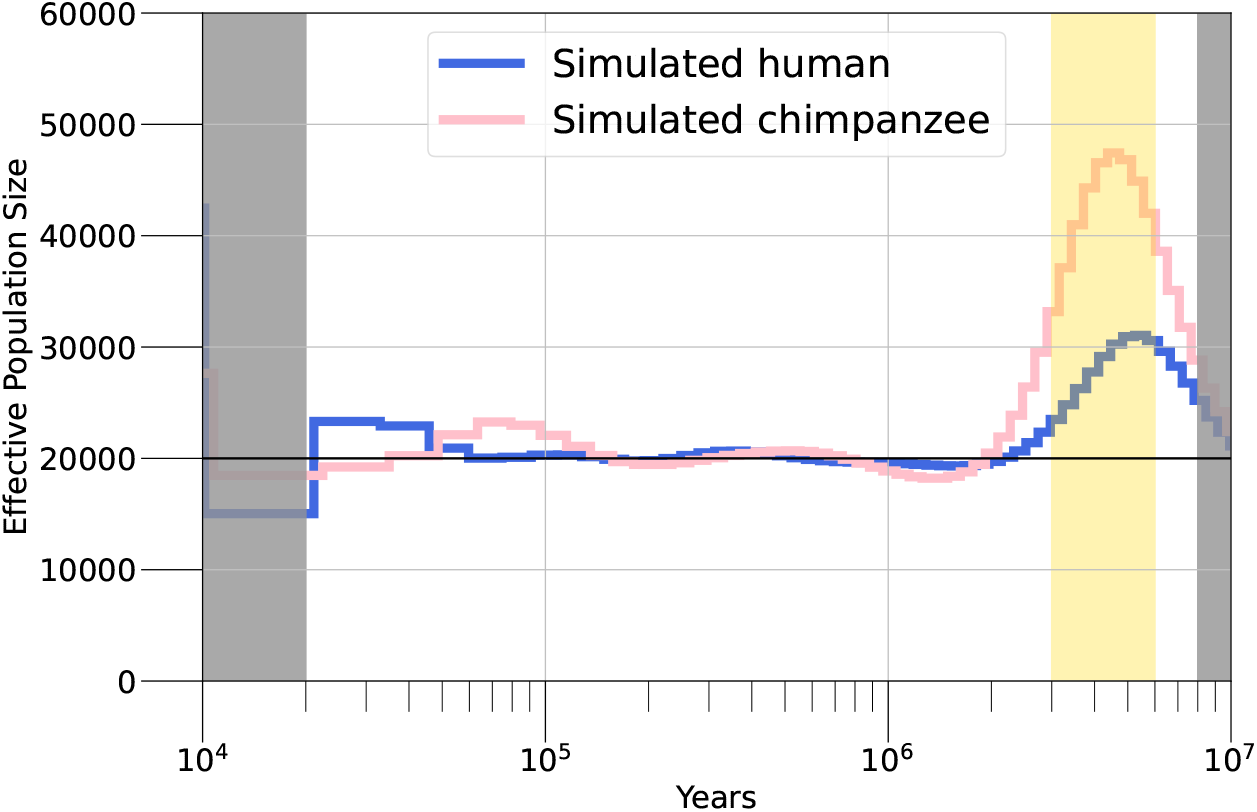
PSMC’ inference on 2Gb of simulated human and chimpanzee sequence. In this simulation, at 6 Mya a panmictic ancestral population of size 20k splits into proto1 and proto2 (each of size 20k). At 3 Mya, human forms as a 80:20 mix of proto1 and proto2, respectively, and chimpanzee forms as a 50:50 mix of proto1 and proto2.

**Figure A3.**
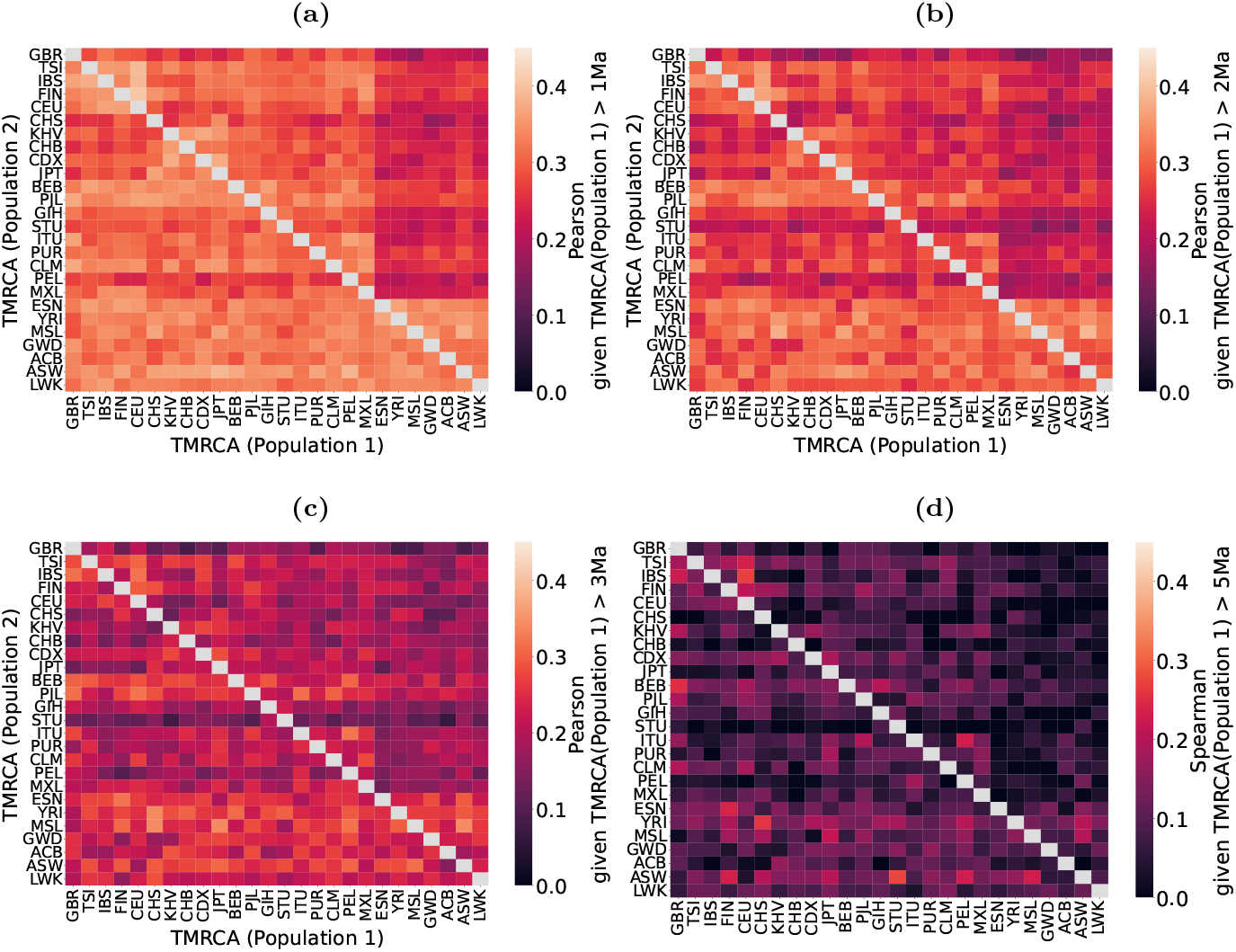
The Pearson correlation between inferred TMRCAs, for all pairwise combinations of two populations from the 1000GP. We calculated these for chromosome 1. To obtain point estimates at each position, we used the posterior mean. For each possible population pair, we look at positions where population 1 (x-axis) is larger than *t* and correlate these with the TMRCAs from population 2 (y-axis; note these are not necessarily larger than *t*, so the matrices are not symmetric). The value of *t* for **(a), (b), (c)**, and **(d)** is 1 Mya, 2 Mya, 3 Mya and 5 Mya, respectively. The average Pearson *r*^2^ for (a), (b), (c), and (d) is 0.3, 0.26, 0.21 and 0.11, respectively, and all correlations have p-value*<*0.05 except for 72 combinations from (d).

**Figure A4.**
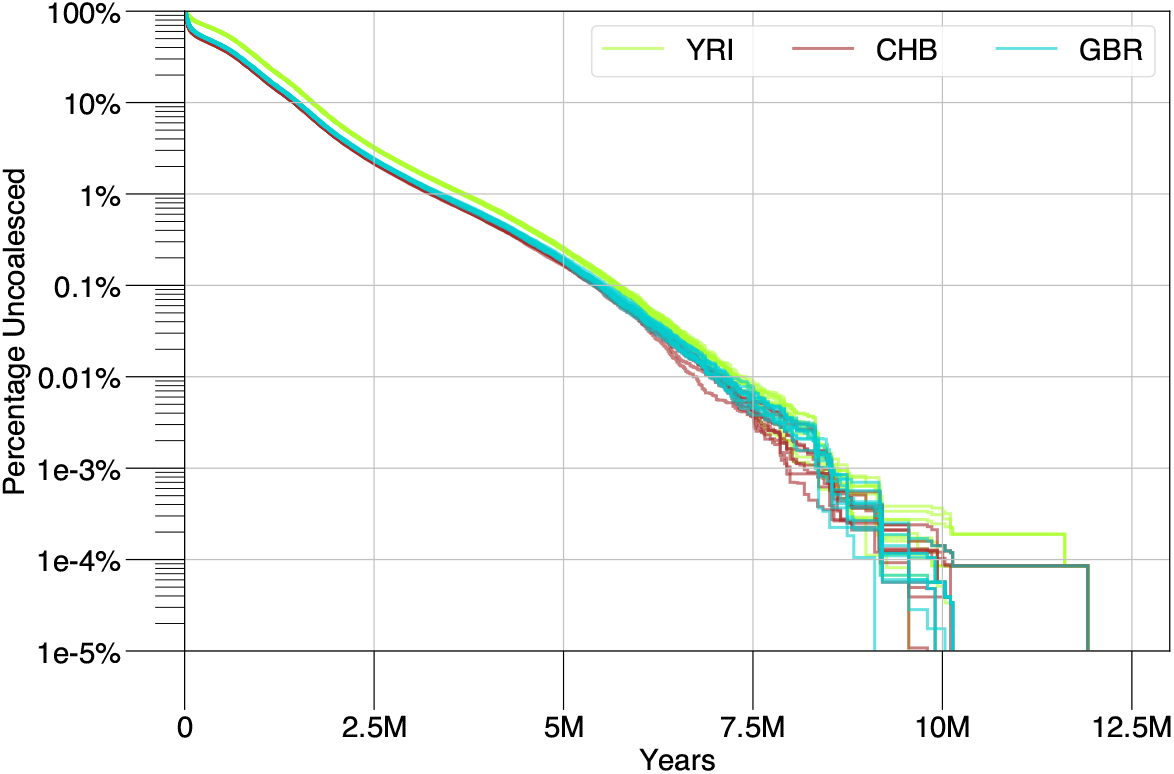
The Relate inferred fraction of uncoalesced genome as a function of time, for 10 diploid samples in the YRI, GBR and CHB populations from 1000GP.

### 5.2 Analysis of PSMC’ model violations

In this section, we analyse the effect of various model misspecifications on the inference from the PSMC’ model. We also analyse how the inferred coalescence times across the genome associate with various annotations, including for example repeat content, distance to coding sequence, strength of background selection, and recombination rate.

#### 5.2.1 Repeats as a cause of artifactual heterozygous signal

As noted in previous work [6], false heterozygotes caused by repeated regions or segmental duplications may lead to excessively long segments with high heterozygosity, which may lead to an excess of ancient coalescent events. To examine whether this might artefactually be creating the observed signal, we analysed the fraction of the genome in repeated regions in the 1000GP data, and stratified this by inferred TMRCA. We observed that the repeat content increases as a function of TMRCA (regression slope ∼ 2%), and that the effect is significant (Pearson’s *r*^2^ ≈ 0.63, p-value*<*0.005; see Methods and Figure A5a). A similar analysis with segmental duplications revealed no significant correlation (Figure A5b). We re-ran our analysis with repeated regions and segmental duplications masked out, and observed that the ancient peak is still present, although when repeats are removed it exhibits less prominence (Figure A6, left and middle panel, respectively). Thus we conclude that repeated regions or segmental duplications are unlikely to be the cause of the ancient peak. We further note that the ancient peak exists in the T2T data for which repeats have been filtered explicitly (see Methods).

#### 5.2.2 Variable mutation rate across the genome

The PSMC’ model as implemented in MSMC2 assumes a constant mutation rate throughout the genome, despite ample evidence that the mutation rate varies significantly by genomic location [98, 99]. Unfortunately, variation in mutation rates across different genomic regions can lead to systematic biases in the estimation of TMRCAs. This is because regions with faster mutation rates will have more mutations than otherwise, which the PSMC’ model will interpret as coming from a more ancient TMRCA, and vice versa. Consequently, overestimation and underestimation of TMRCAs can occur, which can confound efforts to infer accurate evolutionary histories.

**Figure A5.**
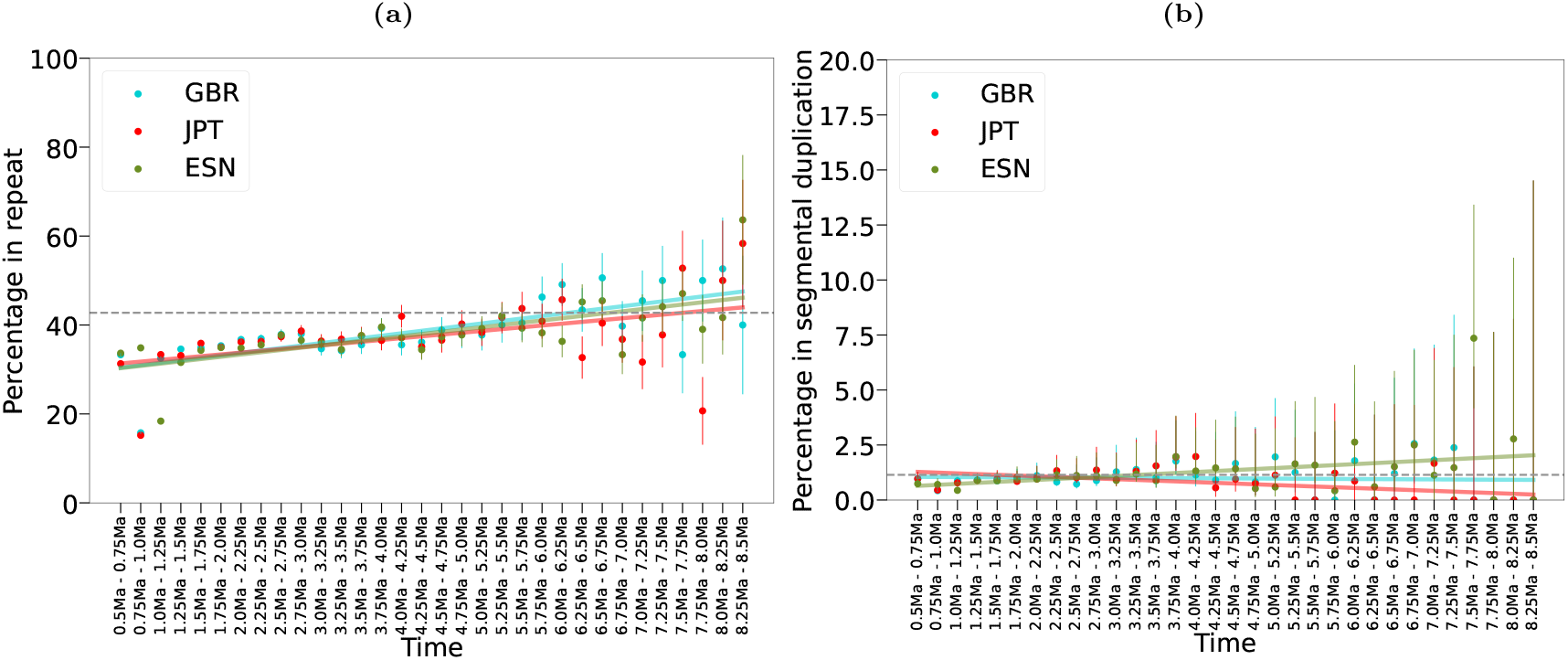
The relationship between the posterior mean TMRCA and the repeat content (a) or segmental duplications (b), on chromosome 1 for one individual each from GBR, JPT, and ESN in the 1000GP. (a) The reported Pearson *r*^2^ for each individual is 0.73, 0.48, 0.69, with p-value*<*0.005 for all. The grey, dashed line indicates the chromosome wide average repeat content. (b) The reported Pearson *r*^2^ for GBR and ESN is not significant, though for JPT it is -0.5 with p-value 0.003.

**Figure A6.**
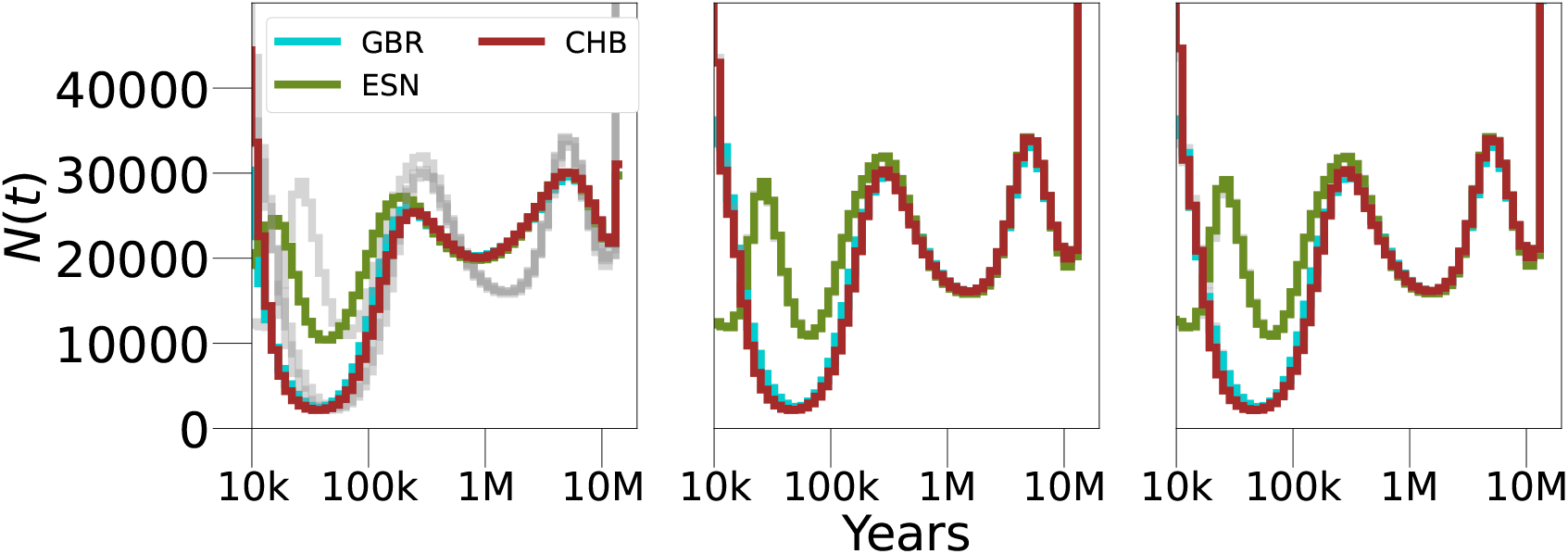
Applying PSMC’ to human samples after masking out repeated regions (a), segmental duplications (b), and CpG islands (c). The original, unmasked results are shown in grey.

To test this effect, we performed a series of simulations of a spatially varying mutation rate that changes according to a gamma distribution, in line with the models proposed in [100]. We generated a mutation rate map as follows. First, we divided the genome into disjoint intervals, whose length is exponentially distributed according to a rate parameter, *CR*. Then, for each interval, we drew a mutation rate from a gamma distribution whose mean is fixed to be 1.25e-8 [90, 91, 92, 93, 94, 95] with a coefficient of variation, *v*. We simulated over *CR* ∈ {10kb, 100kb, 1Mb} and *v* ∈ {0.15,0.25}, covering the range of values of *v* in reference [100]. We simulated diploid genomes according to a coarse piecewise population size trajectory using msprime [97], then generated pointwise mutations according to the mutation rate map. We generated 5 replicates of this simulation and then inferred the population size history using PSMC’. Finally, as a control, we generated an additional 5 replicates using the same procedure, but with a constant mutation rate of 1.25e-8, and infer *N* (*t*) using PSMC’ as well.

**Figure A7.**
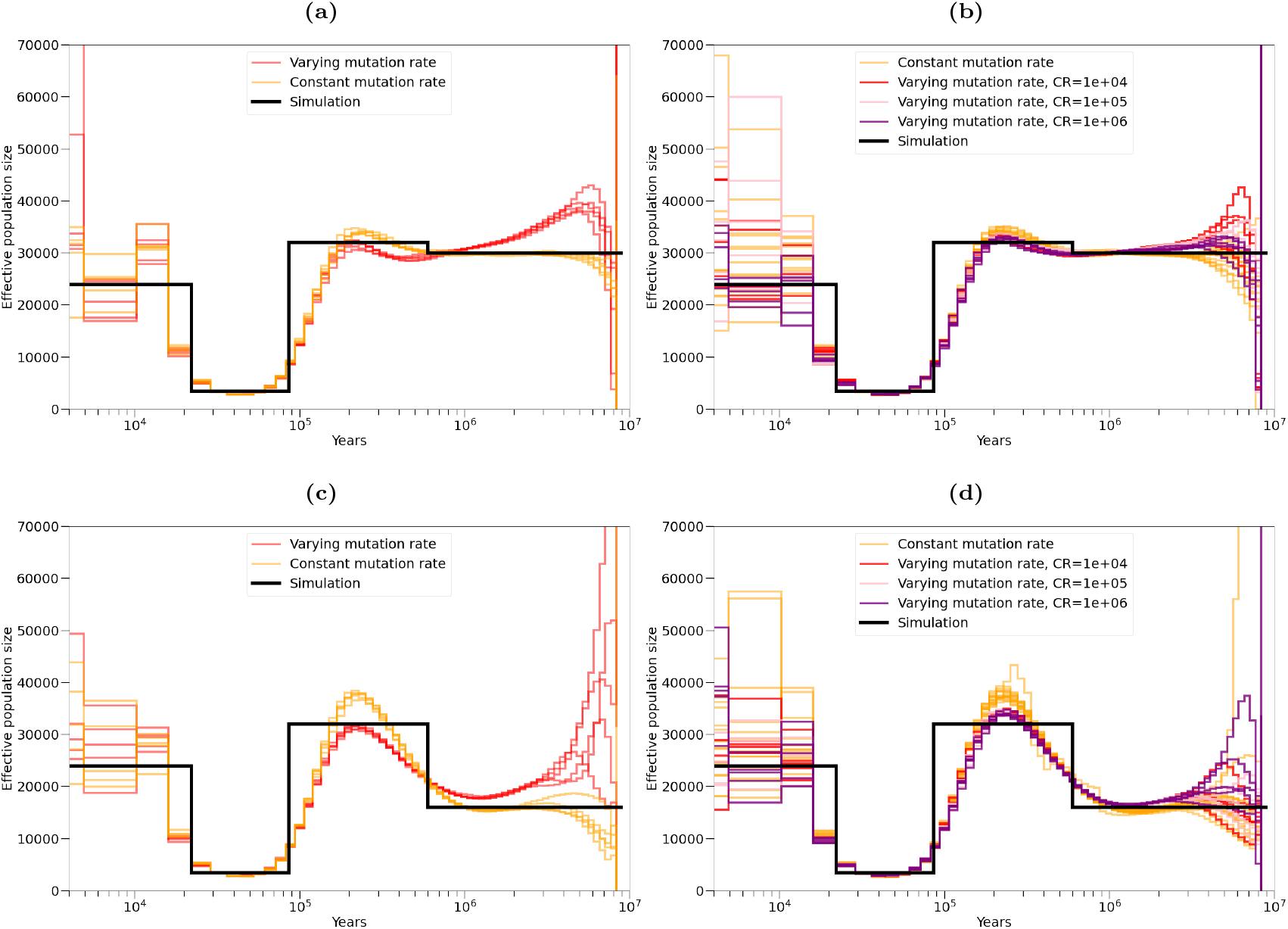
PSMC’ inference on simulations with a mutation rate that varies across the genome. The mutation rate changes at disjoint intervals, whose length is exponentially distributed with rate *CR*. In each interval, the mutation rate is drawn from gamma distribution with mean 1.25e-8 and a coefficient of variation *v*. The red, pink and purple lines show inference from PSMC’ on a simulation where the mutation rate varies, and the gold lines show inference from PSMC’ on a simulation where the mutation rate is constant. **(a)** and **(c)** *v* = 0.25 and *CR* =10kb. **(b)** and **(d)** *v* = 0.15 and *CR*=10kb (red), 100kb (pink), and 1Mb (purple).

This simulation study shows that if the variation in mutation rate is strong enough (*v*=0.15), and the rate of change along the genome is fast enough (*CR*=10kb), PSMC’ detects a spurious peak in ancient time (Figure A7a). However, we note that qualitatively, the false peaks across different replicates vary more in location and height than in the human inference, for which the ancient peaks are well aligned (Figure 1a). Additionally, this simulation used *v*=0.25, which is larger than mutation rate model fitted by [100] for the datasets in [101, 102] (0.18 and 0.15, respectively) but smaller than in [103] (0.27). In a simulation with *v*=0.15, we do not consistently observe a second peak (Figure A7b) for any value of *L*. In a simulation with lower ancient *N* (*t*), *v*=0.25 does generate an artificial inflation of ancient *N* (*t*), but not a discernible peak (Figure A7c). A simulation with *v* = 0.15 and *L*=1Mb again can generate a false peak, though it is not consistent across replicates (purple line, Figure A7d); *CR*=10kb or *CR*=100kb with *v*=0.15 does not seem to have much of an effect as the inference is similar to the constant mutation rate inference.

If the ancient peak was an artefact caused by variable mutation rates, we would expect to see a correlation between the mutation rate and the inferred TMRCA. To test this, we used de-novo mutation abundance counts obtained from trios [104] to generate a pointwise mutation rate map that depends on the local trinucleotide context (see Methods). The generated mutation map is positively correlated with SNP density as reported in 1000GP’s dbSNP (*r*^2^=0.09, p-value*<*1e-100, averaged across non-overlapping windows of 100bp), indicating positions with higher inferred rates have an elevated probability of mutations. We compared the variable mutation rates with the maximum inferred TMRCA across the 26 individuals in 1000GP, evaluated every 1kb. We observed a statistically insignificant Pearson’s correlation (r=0.002, p-value=0.29) and a significant negative Spearman’s correlation (rho=-0.04, p-value=3.7e-93) between TMRCA and mutation rate map. This suggests variable mutation rates do not play a significant role in causing the ancient peak.

Finally, we explored the role of CpG islands in affecting *N* (*t*) inference. CpG islands are regions of DNA characterised by a high frequency of cytosine-guanine dinucleotides. They are often found near the transcription start site of genes and are associated with gene regulation. Due to their regulatory role, CpG islands have fewer mutations than expected by their high GC content, and therefore may contribute to mutation rate variability along the genome. The fraction of the genome in a CpG island is 0.007, and we stratify this by inferred TMRCA in Figure A8. The reported Pearson association between TMRCA and the fraction of regions in a CpG island is negative, though often this is not significant. Moreover in ancient time the CpG fraction seems to increase; to test the effect on inference we masked out CpG islands and inferred *N* (*t*). We still observed an ancient peak (Figure A6, right), and thus concluded that CpG islands are not its cause.

#### 5.2.3 Variable mutation rate through time

The PSMC’ model also assumes a constant mutation rate across past generations. While this is a sufficient approximation to allow inference in the last ∼50 kya [31], it may not hold in more ancient times. Indeed, it has been suggested that the mutation rate has changed over the course of human evolution, also referred to as the “hominoid rate slowdown” [47, 48, 49, 50]. This is based on the observation that the yearly mutation rate estimated from human pedigree studies is almost half the rate inferred by considering observed differences between human and chimpanzee genomes. The effect of a possible slowdown on PSMC’ model estimates is unclear.

To this end, we simulated 10 replicates of a diploid genome, arising from a constant population size, and a mutation rate that is fixed at 2.5e-8 at 10 Mya then decreases to 1.25e-8 at present time (Figure A9a). To simulate a temporally changing mutation rate, we discretised time into *D*=64 segments with 65 time interval boundaries according to equation 1, and set the changes in mutation rate as piecewise constant in these intervals, *µ* = [*µ*_1_, …, *µ*_64_]. We simulate the coalescent process using msprime [97] with Hudson’s model of recombination [105], then utilise the memoryless property of the exponential distribution to add mutations at each position. With a coalescence time of *t* where *τ*_*i*_ ≤ *t < τ*_*i*+1_, measuring time in generations the probability of at least one mutation arising on either lineage is:

**Figure A8.**
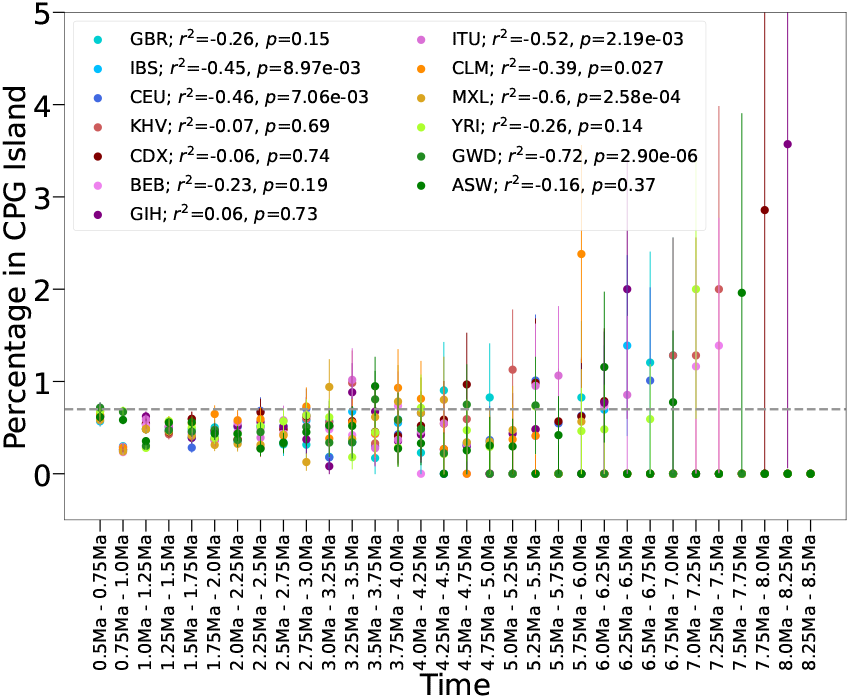
The correlation between the posterior mean TMRCA and the fraction of regions in a CpG island, on chromosome 1 for one individual from GBR, JPT, and ESN in the 1000GP. The Pearson *r*^2^ and p-value are shown in the legend, although the relationship appears to be non-linear, and dominated by noise in ancient TMRCA bins. The dashed line indicates the genome-wide fraction of regions in a CpG island.

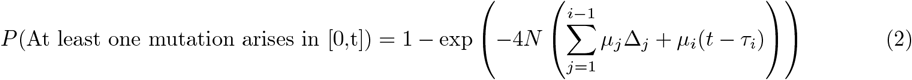

where *i* is the largest integer such that *τ*_*i*_ ≤ *t, τ*_0_ = 0, and Δ_*j*_ = *τ*_*j*_ − *τ*_*j*−1_.

We used PSMC’ to infer an *N* (*t*) curve, and compared the results to *N* (*t*) curves inferred from simulations with a fixed mutation rate of 1.25e-8. We observed that PSMC’ infers an increasingly inflated *N* (*t*) in ancient times (Figure A9b). With the same mutation model, we also simulated changes in *N* (*t*) similar to those inferred in the ESN (Figure 1). Again, we observe the inference of *N* (*t*) from PSMC’ is increasingly inflated in ancient times, though the general shape of the trajectory is recovered (Figure A9c). We note that if the changes in mutation rate are known, we can simply scale the inference appropriately to correct the error (Figure A9d and e). Even though changes in the mutation rate over time can affect the estimation of *N* (*t*), it is unlikely that the observed ancient peak in humans is due to this, as producing a peak as an artefact of changing mutation rates would require several rapid and severe mutation rate fluctuations, which seems unlikely. We conclude that mutation rate variation through time is not a likely cause of an ancient peak, though we note that this may contribute to the differences in ancient *N* (*t*) magnitudes in humans and chimpanzees (Figures 1a and b).

**Figure A9.**
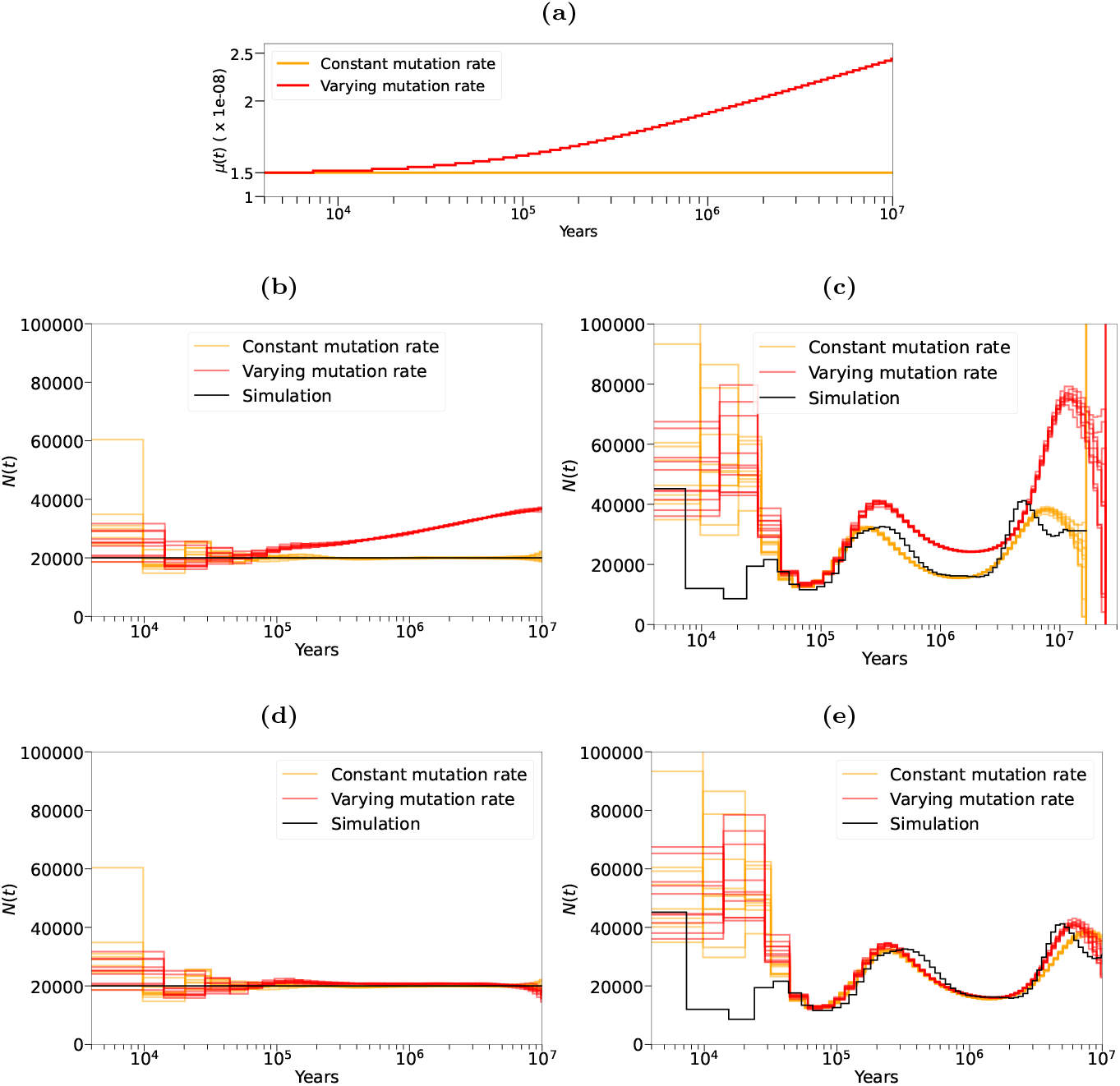
PSMC’ inference on simulations with a mutation rate that varies through time. **(a)** The mutation rate model, *µ*(*t*): at 10 Mya the mutation rate is 2.5e-8, then it slows down to 1.25e-8 at present. **(b)** Inference from PSMC’ on a simulation with constant population size, where the mutation rate changes through time according to the model in (a) in red, and a constant mutation rate (1.25e-8) in gold. **(c)** The same simulation as in (b) but with a population undergoing size changes. **(d)** and **(e)** are the same as (b) and (c), respectively, although the inferred *N* (*t*) on the simulation with a varying mutation rate (red) has been divided through by *µ*(*t*) (red line in (a)).

#### 5.2.4 Variable recombination rate across the genome

The PSMC’ model also assumes a constant recombination rate across the genome, and uniformly through all past generations. However, recombination rate varies along the genome, to a much greater extent than mutation rate, with high rates at recombination hotspots and lower rates in different regions [51, 106, 53, 54, 55]. Looking back in time, the landscape of recombination is known to transform on a time scale of a few hundred thousand years [107, 108, 109, 110]. It is thus unclear how recombination rate variation affects *N* (*t*) inference.

Previous work [6] has demonstrated that *N* (*t*) inference in PSMC is robust even under simulations including recombination hotspots. As additional confirmation, we tested the effect of mis-specifying the recombination rate. Denote the simulated scaled recombination rate as *ρ*; we fix PSMC”s recombination rate 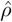 as 0.2*ρ*, 0.5*ρ*, 1*ρ*, 2*ρ*, or 5*ρ*, and infer *N* (*t*) (Figure A10a). We observed that PSMC’ is able to recover *N* (*t*) relatively accurately for all values of the recombination rate except 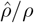 equal to 5. Finally, we saw a significant correlation between the local recombination probability (genetic map taken from HapMap [111, 112]; downloaded from https://alkesgroup.broadinstitute.org/Eagle/) and inferred TMRCAs (Figure A11), though the relationship appears to be non-linear. We investigated this by simulating from this genetic map, and inferring an *N* (*t*) curve assuming a constant recombination rate. We observed that this type of model mis-specification does not significantly alter inference (Figure A10b and A10c). Given these observations, we believe it is unlikely that a spatially varying recombination rate could generate a fake ancient peak.

**Figure A10.**
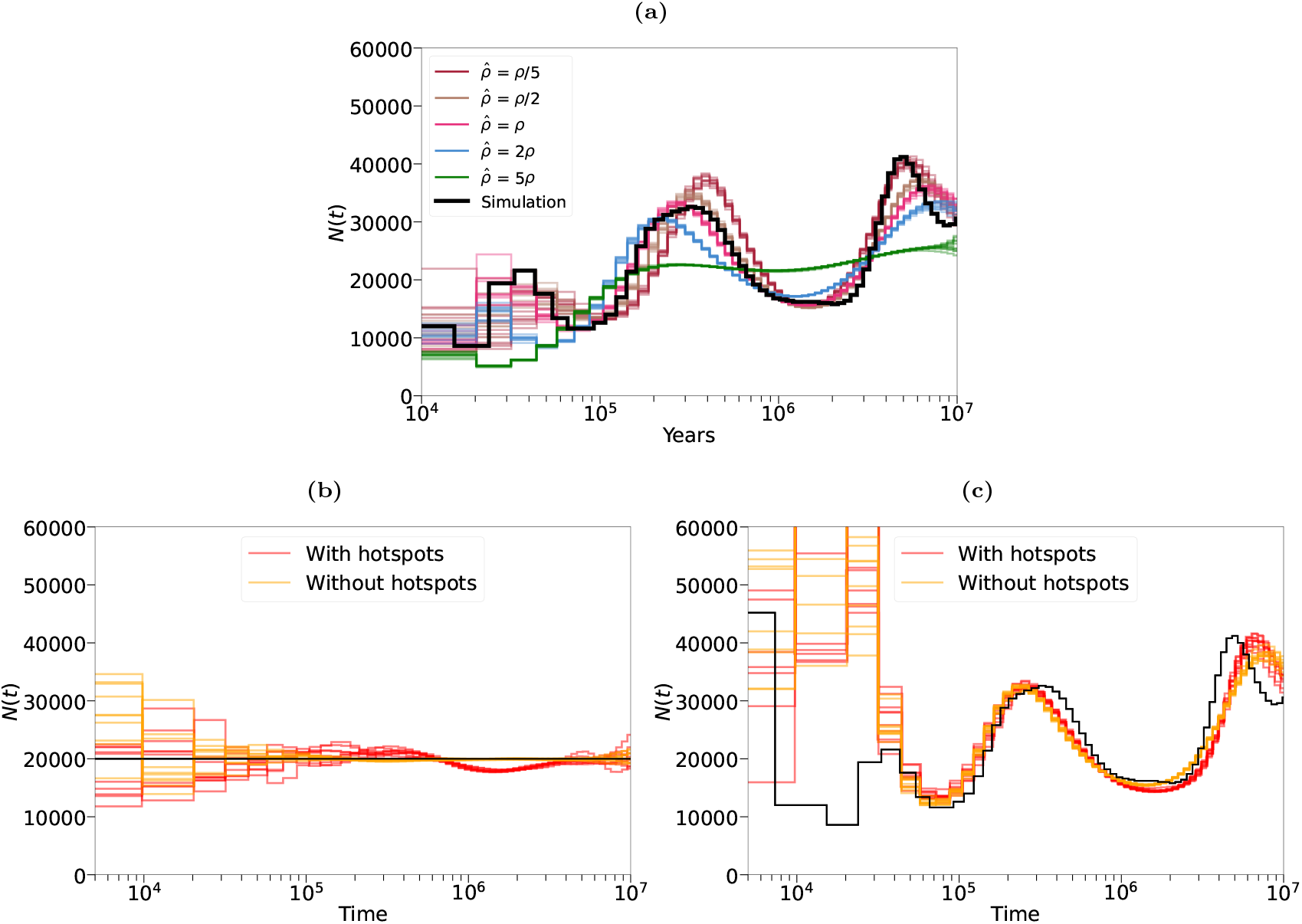
Inference from PSMC’ when its model of recombination is violated. **(a)** In a simulation with constant recombination rate *ρ*, we deliberately fixed the recombination rate used by the PSMC’ algorithm 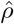 at an incorrect value. The black line indicated the simulated *N* (*t*) and the coloured lines indicate the various values of 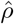, which are expressed relative to the simulated value. **(b)** and **(c)** Inference from PSMC’ on simulations with changes in the recombination rate according to HapMap (red lines) and a constant recombination rate (gold lines).

#### 5.2.5 Recurrent mutations

The PSMC’ model assumes an infinite sites model, in which a site can only experience one mutation. In reality, some ancient sites will have experienced more than one mutation. Under a Jukes-Cantor model of mutation [113], which assumes that each base pair mutates with uniform probability to another, with probability 1/3 a second mutation at the same site will revert to its ancestral state. These will be observed as a homozygote and therefore will tend to lower the estimate of TMRCA by methods based on the PSMC’ model. With probability 2/3 a second mutation at the same site induces a distinct biallelic SNP, which is still observed as a heterozygote and so will not affect TMRCA inference. Overall, repeated mutations at the same site will tend to uniformly and gradually underestimate TMRCA with increasing time, but the effect will be very small given hominin heterozygosity, of the order of 1% out to 10 million years.

**Figure A11.**
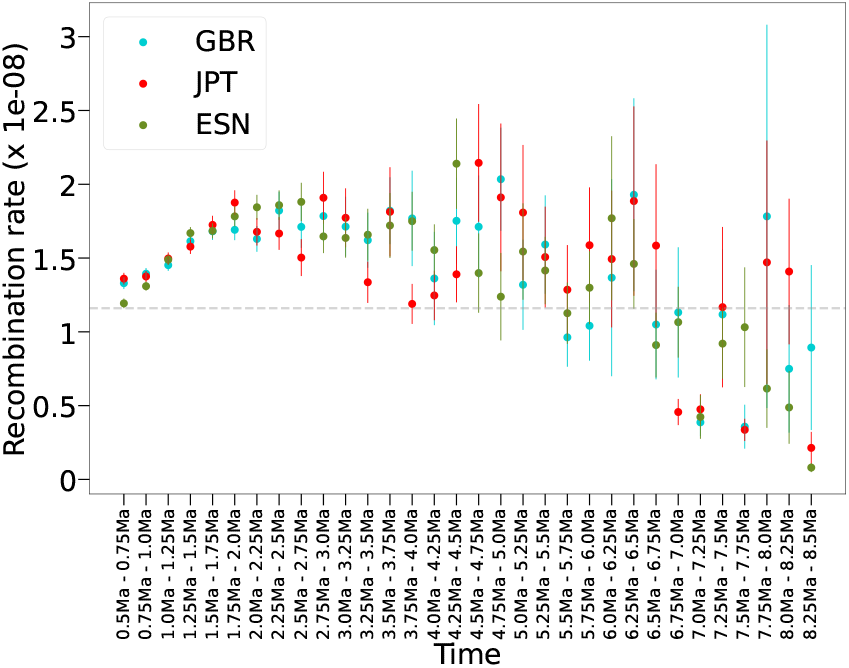
The relationship between the posterior mean TMRCA and the HapMap recombination rate map, on chromosome 1 for one individual from GBR, JPT, and ESN in the 1000GP. The reported Pearson *r*^2^ for each population is 0.45, 0.41, and 0.50, with p-value*<*0.02 for each. However, clearly the relationship is non-linear; it appears that recombination rate increases as does TMRCA up until ∼ 2 Mya, after which they become negatively correlated. This may come from model violations, in that the PSMC model underlying PSMC’ assumes a uniform rate of recombination, or that the recombination landscape changes over time [107]. The grey, dashed line indicates the chromosome wide average recombination rate.

Indeed, in almost all the simulations previously presented a Jukes-Cantor model [113] was used to generate mutations, in which recurrent mutations are allowed to occur. Generally, in simulations with uniform *µ* in time and space, we observe that inference of ancient *N* (*t*) is reasonably accurate in ancient time (Figures A7, A9, and A10).

#### 5.2.6 Background selection

Background selection (BGS) is a form of linked selection, where the removal of deleterious mutations reduces genetic diversity in the surrounding regions due to linkage [114, 115]. It has been demonstrated that BGS is pervasive throughout the human genome, and that this explains roughly 60% of the variance in diversity at the megabase scale [56, 57]. Most demographic inference methods, however, including PSMC’, assume that the genome evolves neutrally. This is problematic, as it has been shown that wrongly assuming the absence of selection means that inference methods are not able to accurately reconstruct the demographic history [58, 60, 59].

We correlated the inferred TMRCAs against a high resolution map that describes the strength of BGS across the human genome [57]. This map infers a local B value, which is a measure of the reduction in local diversity due to BGS and so inversely related to the strength of BGS. The B value is calculated by integrating functional and genetic map information. We saw a significant positive correlation (Pearson *r*^2^ ≈ 0.7, p-value*<*1e-5; Figure A12) between B value and TMRCA, which is consistent with expectations that reduced diversity will be correlated with reduced coalescent time [115]. To explore the effect of BGS on inference of *N* (*t*) in real data, the authors in [62] binned the genome of 10 YRI individuals into quintiles according the strength of BGS and applied PSMC’ to each (reproduced in Figure A13). In all quintiles except the one with strongest BGS, a clear second peak is seen in ancient time. The peak is not seen in the strongest BGS quintile likely because all of the input sequence has already coalesced. In none of the simulations with realistic parameters of BGS, as shown in [59, 116, 62, 117], does PSMC’ infer a false ancient peak. We thus find it implausible that widespread BGS could generate an ancient peak as observed in humans.

**Figure A12.**
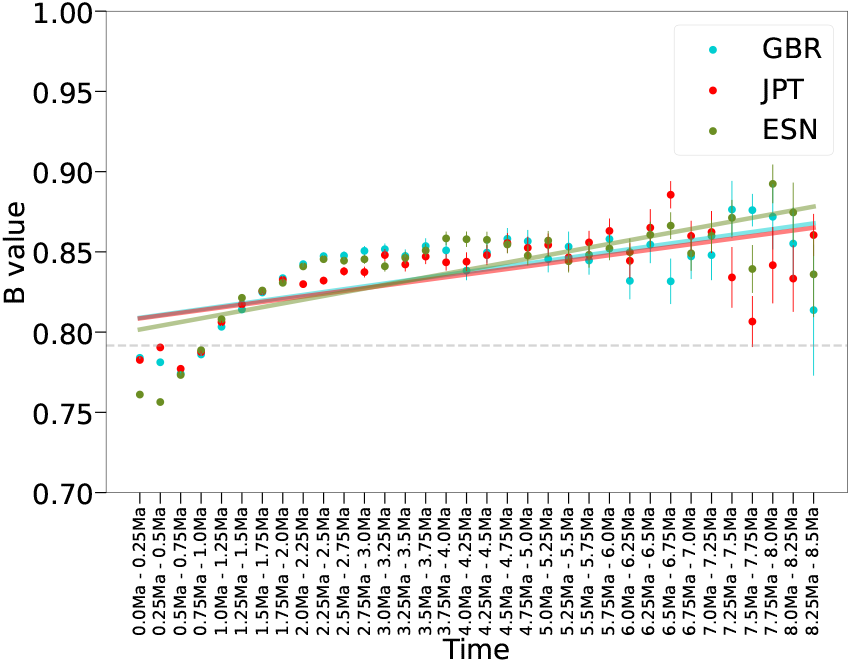
The relationship between the posterior mean TMRCA and *B*, which is a high resolution map of the effects of background selection [57], on chromosome 1 for one individual from GBR, JPT, and ESN in the 1000GP. *B* values indicate the local strength of BGS, with 1 being no reduction in genetic diversity due to selection and 0 being full removal of genetic diversity. The reported Pearson *r*^2^ for each population is 0.67, 0.67, and 0.75, with p-value*<*1.3e-5 for each. In general this suggests that B value increases with TMRCA (a linear line of best fit has been added to show this), though the relationship appears to be non-linear. The grey, dashed line indicates the chromosome wide average B value.

**Figure A13.**
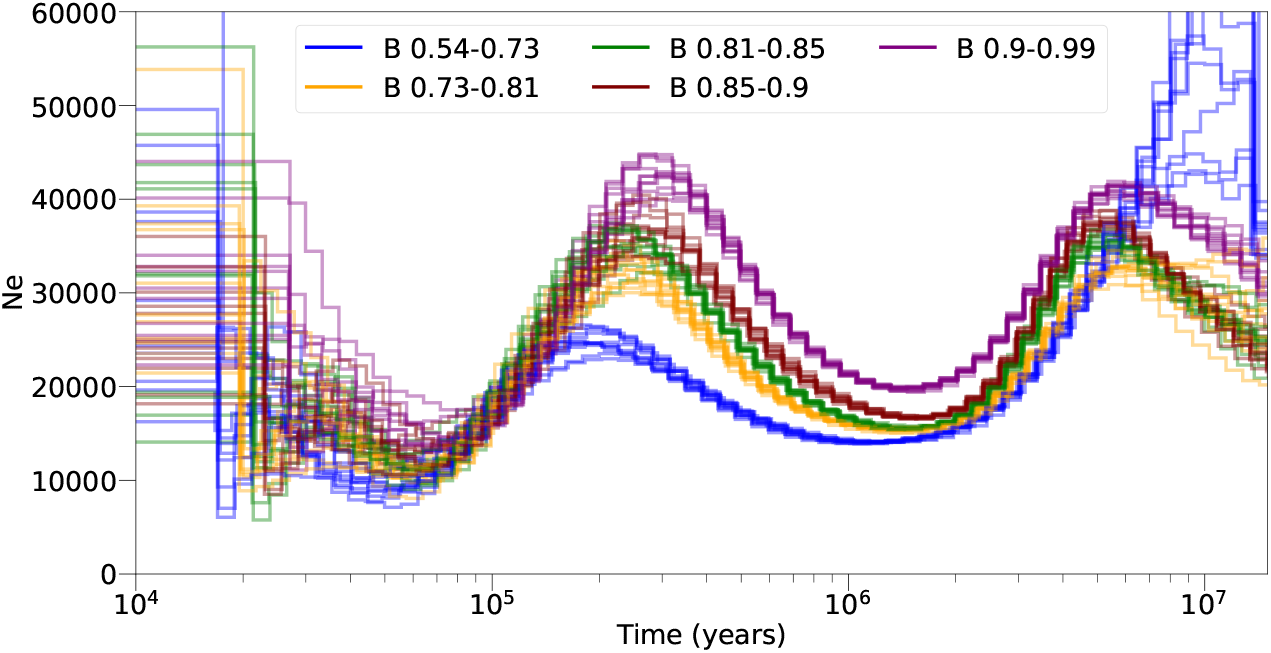
PSMC’ inference of *N* (*t*) in 10 YRI individuals from the 1000GP project, stratified into quintiles according to the strength of BGS (as inferred by [57]). This figure is reproduced with permission from [62].

**Figure A14.**
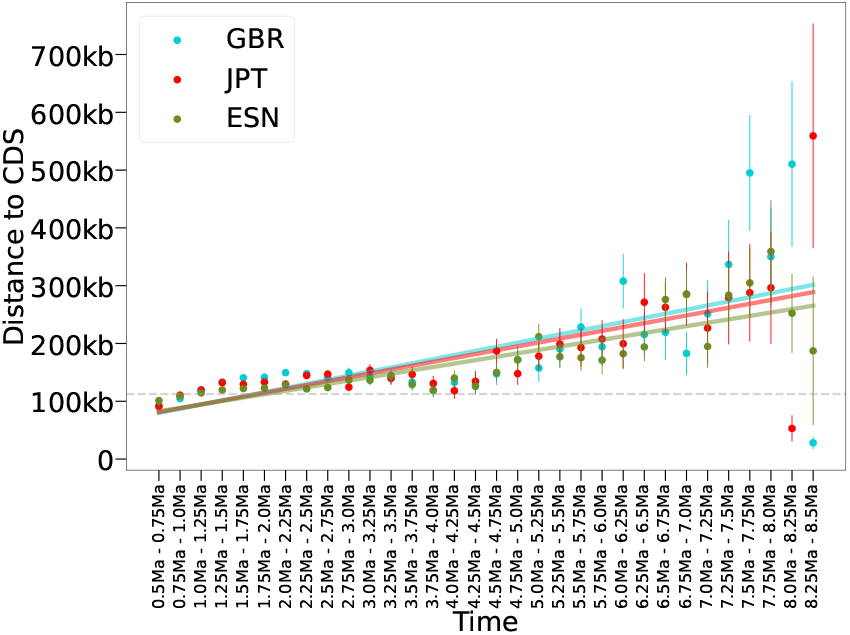
The relationship between the posterior mean TMRCA and mean distance to coding sequence (CDS), on chromosome 1 for one individual from GBR, JPT, and ESN in the 1000GP. The reported Pearson *r*^2^ for each population is 0.63, 0.67, and 0.83, with p-value*<*0.0001 for each. In general this suggests that dcCDS increases with TMRCA (a linear line of best fit has been added to show this), although the relationship appears reversed from ∼ 3 Mya to ∼ 4.5Ma. The grey, dashed line indicates the chromosome wide average distance to CDS.

#### 5.2.7 Balancing selection

Balancing selection (BLS) is a type of natural selection that maintains genetic diversity in a population by favouring the maintenance of multiple alleles. This type of selection can occur through various mechanisms, such as heterozygote advantage, frequency-dependent selection, or spatially varying selection [61]. BLS operating for a long time period will maintain advantageous polymorphism and result in an older TMRCA than expected under neutrality. Therefore, if BLS were sufficiently prevalent in the genome, this would manifest as enrichment of older coalescence times, which would increase inferred *N* (*t*) in the past.

The prevalence of BLS in the human genome is unclear. Initially considered a rarity [118, 119] and overlooked, balanced polymorphisms have recently received renewed attention with several lines of evidence showing their relevance in human evolution [120, 121, 122]. In one study, hundreds of loci were implicated in possible trans-species BLS, maintained since earlier than human-chimpanzee speciation [123]. By finding regions of ancient shared ancestry across 54 individuals from Complete Genomics [124], in [125] the authors suggest numerous regions that are under BLS. We calculated the correlation between inferred TMRCAs and distance to coding sequence (CDS), and found that ancient TMRCAs tend to be increasingly far away from CDS (Figure A14; Pearson *r*^2^ ≈ 0.71, p-value*<*0.0001). As BLS usually acts on or near functionally important parts of the genome, this makes it unlikely that the ancient TMRCAs underlying our ancient peak are driven by BLS and that this is the cause of the ancient peak.

### 5.3 Methods

#### 5.3.1 TMRCA Correlations with annotations

We downloaded the the tracks for repeats, segmental duplications, and CpG islands from the UCSC Genome Browser https://genome.ucsc.edu/cgi-bin/hgTables [126]. The positions of coding sequence were obtained from Gencode ftp.ebi.ac.uk/pub/databases/gencode/Gencode_human/release_45/gencode.v45.chr_patch_hapl_scaff.basic.annotation.gff3.gz. We realigned all 1000 Genomes Project data to GRCh38 for comparison with various annotations. We used the B-map as inferred by Murphy et al. [57] and lifted over from GRCh37 to GRCh38 [126].

We took the posterior mean TMRCA across chromosome 1 for the 26 1000GP samples. We stratified the TMRCAs into windows of 250 kya and analysed how various functional annotations correspond to these TMRCA windows, and ensured that positions in the analysis passed the strict mappability mask.

To generate a pointwise mutation rate map, we used strand-symmetric de-novo abundance counts obtained from trios [104] to calculate the relative frequencies of each possible mutation in its trinucleotide context. To get a pointwise mutation rate per trinucleotide, we scaled these frequencies appropriately to obtain the genomewide mutation rate of 1.25e-8.

